# Giving you five: A neuroimaging atlas of the nigrosomes in the substantia nigra based on 3D histology

**DOI:** 10.64898/2026.04.24.720555

**Authors:** Malte Brammerloh, Anneke Alkemade, Pierre-Louis Bazin, Caroline Jantzen, Carsten Jäger, Andreas Herrler, Kerrin J. Pine, Markus Morawski, Rawien Balesar, Katrin Amunts, Birte U. Forstmann, Nikolaus Weiskopf, Evgeniya Kirilina

## Abstract

Nigrosomes are formed by clusters of pigmented dopaminergic cells in the *substantia nigra* that critically contribute to dopaminergic function. The ever-increasing resolution of ultra-high-field MRI brings clinical imaging of these clusters into reach, promising unprecedented insight into the functional role of the nigrosomes and their early degeneration in Parkinson’s disease. However, due to the nigrosomes’ small extents and intricate shapes, they are not included in current MRI brain atlases, preventing nigrosome-specific MRI data analysis. We provide a comprehensive 3D histological atlas of the five nigrosomes co-aligned to the widely-used MNI152 2009b space. This atlas is based on 3D-reconstructed, ultra-high-resolution block-face images and gold-standard nigrosome delineations in calbindin-D28K immunohistochemistry. We validated the atlas’s accuracy using the multimodal ultra-high-resolution *post mortem* BigBrain dataset and demonstrated its consistency with qualitative nigrosome atlases based on classical 2D histology. We provide detailed usage instructions for applying our atlas to ultra-high-resolution and -field MRI data. The openly available atlas enables neuroimaging studies of the nigrosomes, opening a new avenue toward understanding the differential involvement of the nigrosomes in the healthy and diseased brain and the development of neuroimaging biomarkers of dopaminergic neurodegeneration.

## Background & Summary

The kidney-bean-sized *substantia nigra* (SN) is a nucleus in the midbrain, situated between the medio-rostral *nucleus ruber* (NR), and the ventro-lateral *crus cerebri*^1,2^, with a wide-ranging impact on brain function (Fig. 1). Classical neuroanatomical studies identified three distinct anatomical layers (tiers) within SN, running parallel to its shortest dimension. The most lateral SN layer, called *pars reticulata*, contains GABAergic neurons within the SN. Two dorso-medial layers, referred to as the ventral and dorsal tiers, belong to the *pars compacta*. They contain neuromelanin-pigmented dopaminergic neurons (DNs), which synthesize most cortical-acting dopamine and contribute to key cognitive functions, including reward-based learning and motor control. Neuromelanin-pigmented dopaminergic neurons form five neuron-dense clusters, called nigrosomes^3^, with submillimeter volumes, and which show low calbindin-D28K immunoreactivity^3^ (Fig. 1A). The middle layer (ventral tier) contains the largest nigrosome 1 (N1), whereas the upper layer (dorsal tier) hosts nigrosomes 2-5 (N2-5) (Fig. 1B). Nigrosomes likely differ in their functional specializations, as was extrapolated from non-human primate studies^4^. Remarkably, in Parkinson’s disease, the five nigrosomes exhibit different spatio-temporal patterns of neurodegeneration^5^ (Fig. 1C). The loss of dopaminergic neurons in Parkinson’s disease is earliest and most pronounced in N1. This loss implies a lack of striatal dopamine and ultimately major motor symptoms. Regarding their functional and pathophysiological importance, mapping nigrosomes opens a unique window into the dopaminergic system.

**Figure 1.**
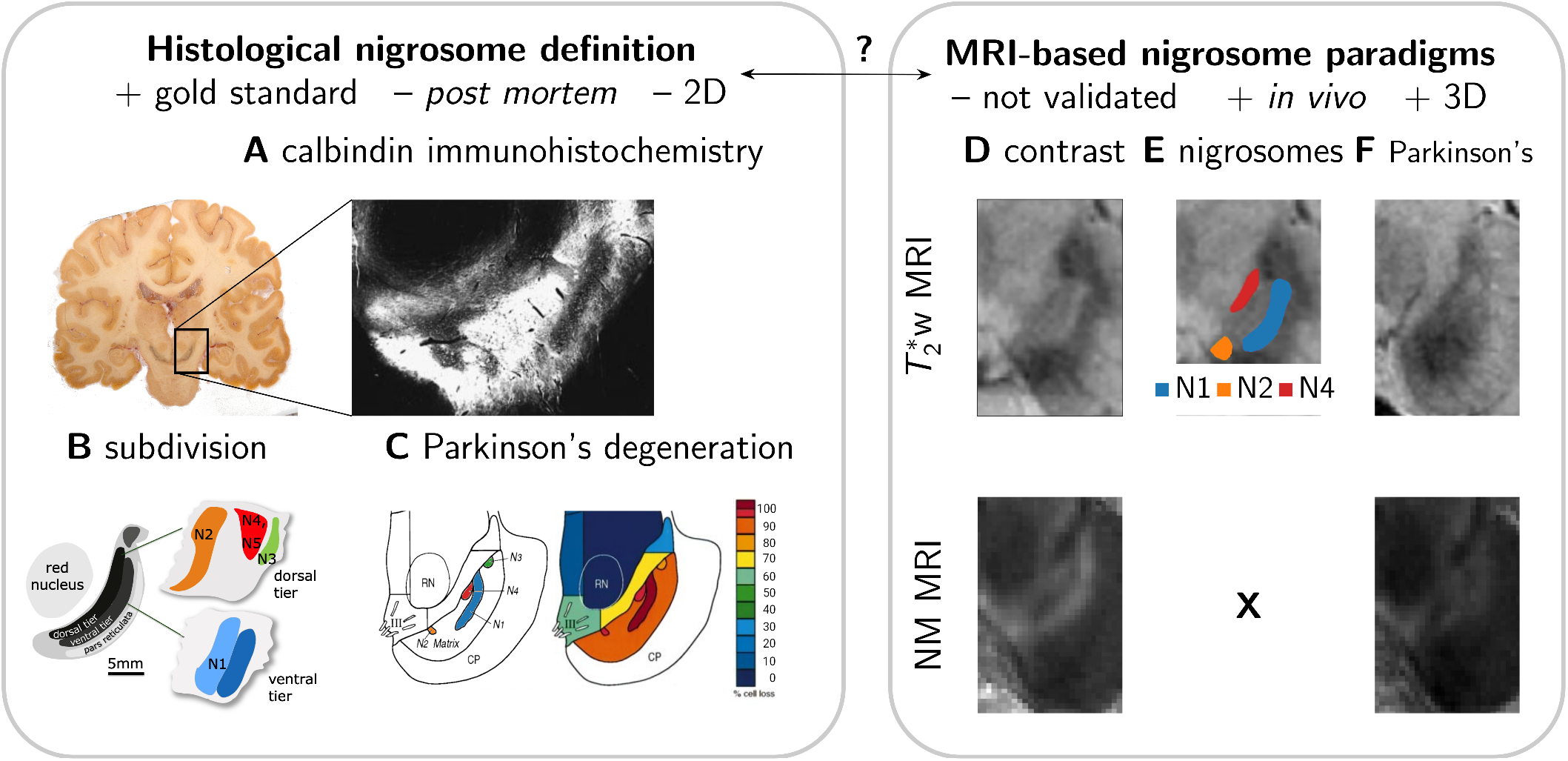
The relationship between nigrosome definitions in histology (A-C) and MRI (D-F) is unclear, rendering nigrosome MRI unfeasible. A: The gold standard nigrosome delineation is based on anti-calbindin-D28K immunohistochemistry, in which nigrosomes appear as areas of reduced immunoreactivity within the highly immunoreactive SN^3^. B: Nigrosomes N1-N5 are defined as areas of low immunoreactivity with consistent spatial arrangement within the dorsal and ventral tiers of the *substantia nigra* (SN) *pars compacta*. The schematic was adapted from^2^. C: Selective degeneration of dopaminergic neurons in the nigrosomes is the hallmark of Parkinson’s disease, with the most severe loss affecting the largest nigrosome N1^5^. On the left is the original delineation of the nigrosomes in an oblique axial section performed by Damier *et al*.^3^. D: The two most prominent MRI techniques to image the nigrosomes are 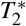-weighted MRI (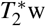 MRI, top row) and an MRI sequence sensitive to neuromelanin, the predominant iron-binding form in dopaminergic neurons (NM MRI, bottom row). Based on the image contrast, a delineation of nigrosomes directly in 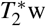 MR images has been proposed^9^, while such an anatomical characterization is currently unavailable for NM MRI. E: MRI research proposed that the nigrosomes correspond to hyperintense areas in 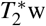 MR images, surrounded by the hypointense SN^9^. However, a recent study questioned this approach to delineate nigrosomes^10^, In NM MRI, a hyperintense stripe within the SN has been observed^11^, possibly corresponding to the stripe-like nigrosome 1^3^. F: Both MRI modalities detect alterations in Parkinson’s disease patients: A loss of hyperintense structures in 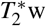 MRI^9^ and, similarly, a loss of the increased contrast-to-noise ratio in NM MRI^11^. The relationship of these alterations to the neuronal loss in the nigrosomes of Parkinson’s disease patients (B) is currently unclear, as no independent validation of the MRI-based nigrosome delineation has been possible.

Recent advances in ultra-high-field magnetic resonance imaging (MRI) at sub-millimeter resolution^6^ bring non-invasive *in vivo* imaging of the nigrosomes into reach. Past research leveraged mainly two MRI techniques for nigrosome imaging: 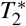-weighted (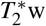) imaging, sensitive to iron in DNs, and a sequence sensitive to neuromelanin, the most abundant iron storage polymer in DNs^7^ (Figs. 1D-1F). Early studies of 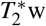 images identified hyperintense areas as nigrosomes, although only the identification of nigrosome 1 was informed by histochemistry^8,9^ (Figs. 1D-2F, top row). Moreover, the identification of hyperintense areas in 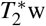 images with the nigrosomes was recently challenged by comparing histochemistry and MRI^10^. Recent progress in neuromelanin-sensitive MRI at 7 T allowed visualization of rich contrast features within the SN, which may reflect the nigrosome anatomy^11^(Figs. 1D-1F, bottom row). However, the relationship between these features and histologically defined nigrosomes is currently unclear. Systematic validation of nigrosome neuroimaging is hampered by the lack of histology-based nigral 3D atlases.

Direct imaging of nigrosomes is complicated by their small extent, comparable with the resolution of state-of-the-art ultra-high-field MRI (approximately 0.5 mm). Neuroimaging atlases that integrate neuroimaging and histochemistry support imaging of anatomical structures with unclear delineation on MR images. Current progress in subcortical atlases has allowed imaging of various brainstem structures, including small nuclei^12–18^. However, none of the available neuroimaging atlases includes the nigrosomes, as three major challenges have remained unresolved. First, an accurate three-dimensional (3D) delineation of the nigrosomes is indispensable. While calbindin immunohistochemistry provides an accurate delineation of the nigrosomes^3^, histological sections suffer from non-linear deformation due to tissue sectioning and histochemical processing, rendering a registration with high enough accuracy for nigrosome alignment challenging. Second, the 3D nigrosome segmentations from several brains need to be aligned to assess the intersubject variability of the nigrosomes. Hence, this registration has to be precise enough to align sub-millimeter-sized structures, while not relying on the contrast of the structures themselves, as otherwise biological variability would be underrepresented in the atlas. Third, dedicated registration procedures are needed for aligning the nigrosome atlas to MRI data. State-of-the-art subcortical atlases achieve a registration accuracy of about 0.5 mm^6,12,19^, hence an accuracy at which the similarly sized nigrosomes could be easily misaligned after registration^12^. As a prerequisite, the atlas should be registered with high precision to an ultra-high-resolution MRI template.

Addressing these challenges, we present a histological nigrosome atlas of SN substructures for MRI, combining 3D-reconstructed block-face images (BFI) and immunohistochemistry (Fig. 2). We created ultra-high-resolution 3D segmentations of the nigrosomes and the SN in BFI. We validated them against the gold-standard definitions using calbindin-D28K immunohistochemistry (Figs. 2A, 2B). We precisely co-registered these segmentations to the AHEAD template^6^ in the MNI152 2009b space^20^, using the SN segmentation for registration. This enabled us to create a nigrosome atlas in MNI152 2009b space (Fig. 2C). We present coregistration procedures for aligning the nigrosome atlas to ultra-high-resolution MRI data and validate their accuracy on the nigrosome length scale using a newly acquired multi-modal BigBrain dataset^21^ (Fig. 2D). We present a use case of our atlas for ultra-high-resolution MRI data acquired at 7 T and provide Python scripts that enable seamless integration of our atlas into state-of-the-art MRI data analysis (Fig. 2E).

**Figure 2.**
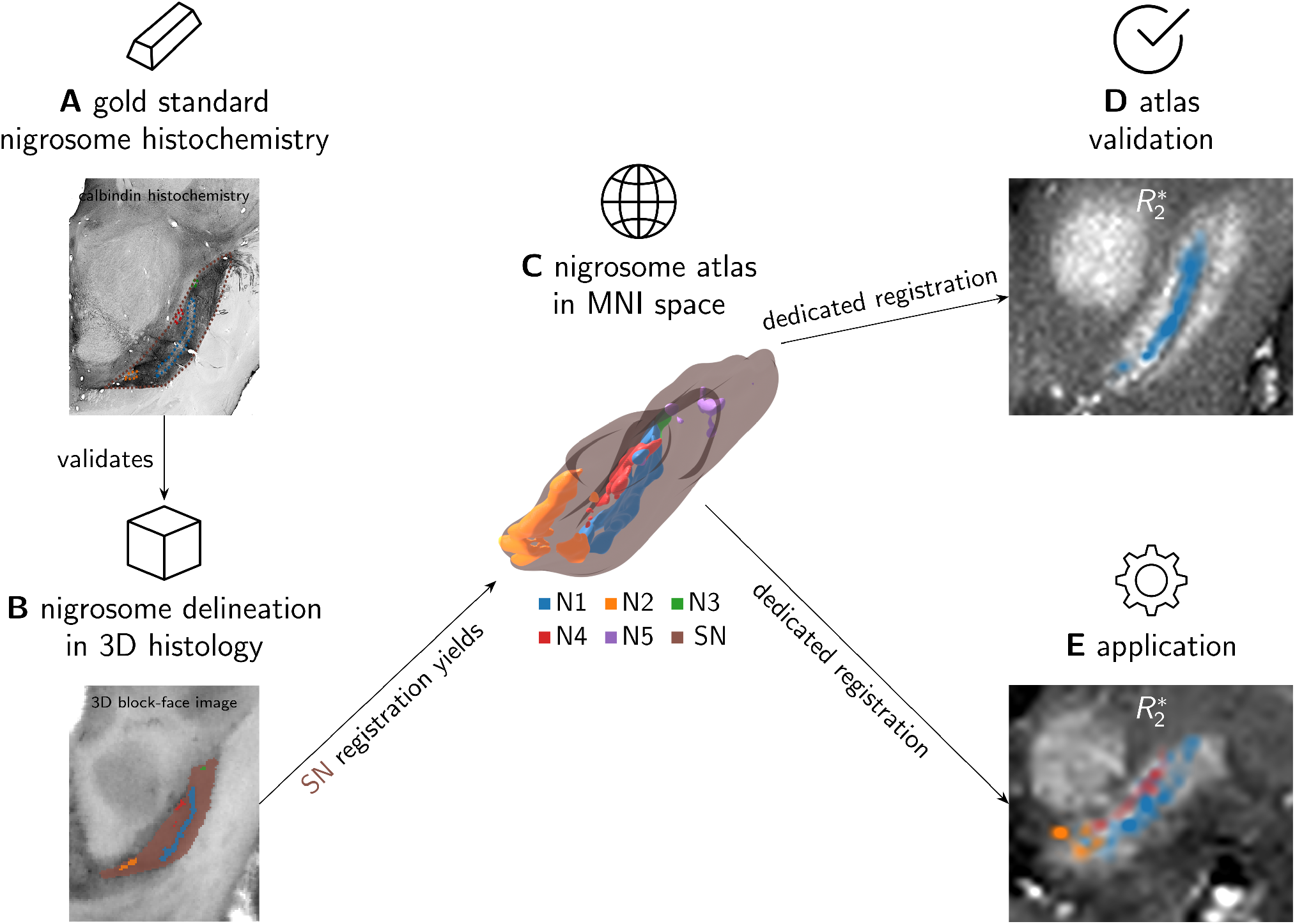
Creating, validating, and applying the nigrosome atlas. We used histological gold-standard definitions of the nigrosomes and the *substantia nigra* (A, gold bar icon) to validate the 3D ultra-high-resolution delineation of these structures (B, cube icon). The icons are used in the subsequent figures to indicate to which step the figure belongs. A precise co-registration of the 3D ultra-high-resolution datasets to the AHEAD template^6^ formed the basis of an MRI-compatible nigrosome atlas (C, map icon), available in the space of the widely-used MNI152 2009b template^20^. To validate the atlas (D, tick icon), we applied it to the multi-modal ultra-high-resolution BigBrain dataset. To this end, we compared the location of nigrosome 1 according to our atlas to its previously reported contrast in 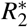 maps acquired *post mortem*^43^. As an application example (E, gear icon), we applied the nigrosome atlas to an *in vivo* MRI dataset acquired with submillimeter-resolution at 7 T. Throughout the manuscript, the icons link figures and subfigures to the conceptual steps illustrated in this figure. Icons made by Smashicons, Darius Dan, srip, Pixel perfect, and DinosoftLabs from www.flaticon.com.

## Methods

### *Post mortem* tissue processing

We used previously published block-face images and anti-calbindin immunohistochemistry for 3D ultra-high-resolution delineations and gold-standard definitions of nigrosomes and the SN^19,22^. In the following, we summarize the tissue processing, while for a detailed description, we refer to the original publications^19,22^. Four previously described *post mortem* specimens (specimens 1, 6, 7, and 8, see Table 1 in^22^) were histologically processed [(73 ± 10) years old, 2 male] (Table 1). The tissue specimens were sourced through the whole-body donation program at the University of Maastricht, the Netherlands, with written informed consent for whole-body donation before the donors’ death and prepared as described before^22^. No donor had a clinical record of neurological disease. Additional approval from the Ethics Committee of the Medical Faculty of the University of Leipzig, Germany (153/17-ek), was obtained for the MRI scans in Leipzig. Tissue was perfusion-fixed within 24 h after the donor’s demise with a formaldehyde-ethanol mixture followed by postfixation for 30 days.

**Table 1.**
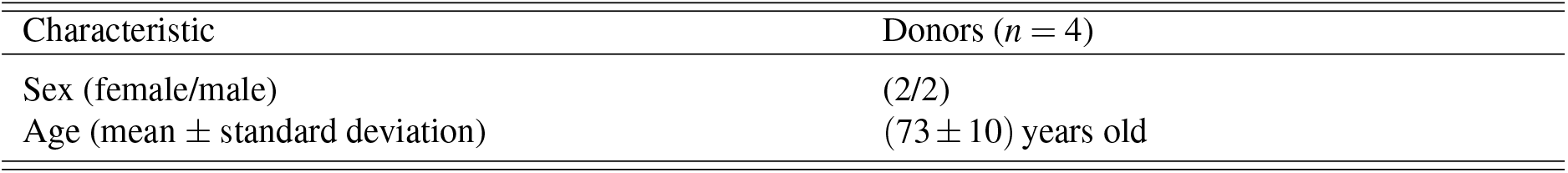
Demographic characteristic of whole-body donors.

### Tissue cutting and 3D block-face imaging

We obtained 3D block-face images (BFI) with a resolution of 150 × 150 × 200 µm^3^ as described in detail before^19,22^ (Figs. 3A-3C). In short, brains were extracted, saturated in sucrose, slowly frozen, and embedded in TissueTek OCT resin before they were cut into 200 µm thick coronal sections using a cryomacrotome. Block-face images were acquired before cutting each section, achieving an isotropic in-plane resolution of (150 µm)^2^. One tissue specimen broke during cutting, but the *substantia nigra* remained intact, allowing us to use it for segmenting the nigrosomes. 3D BFI were reconstructed by stacking BFI without further alignment and converting BFI from color to a gray-scale lightness index. To improve the visibility of nigrosomes, an edge-preserving median image filter was applied to the 3D BF^22^.

**Figure 3.**
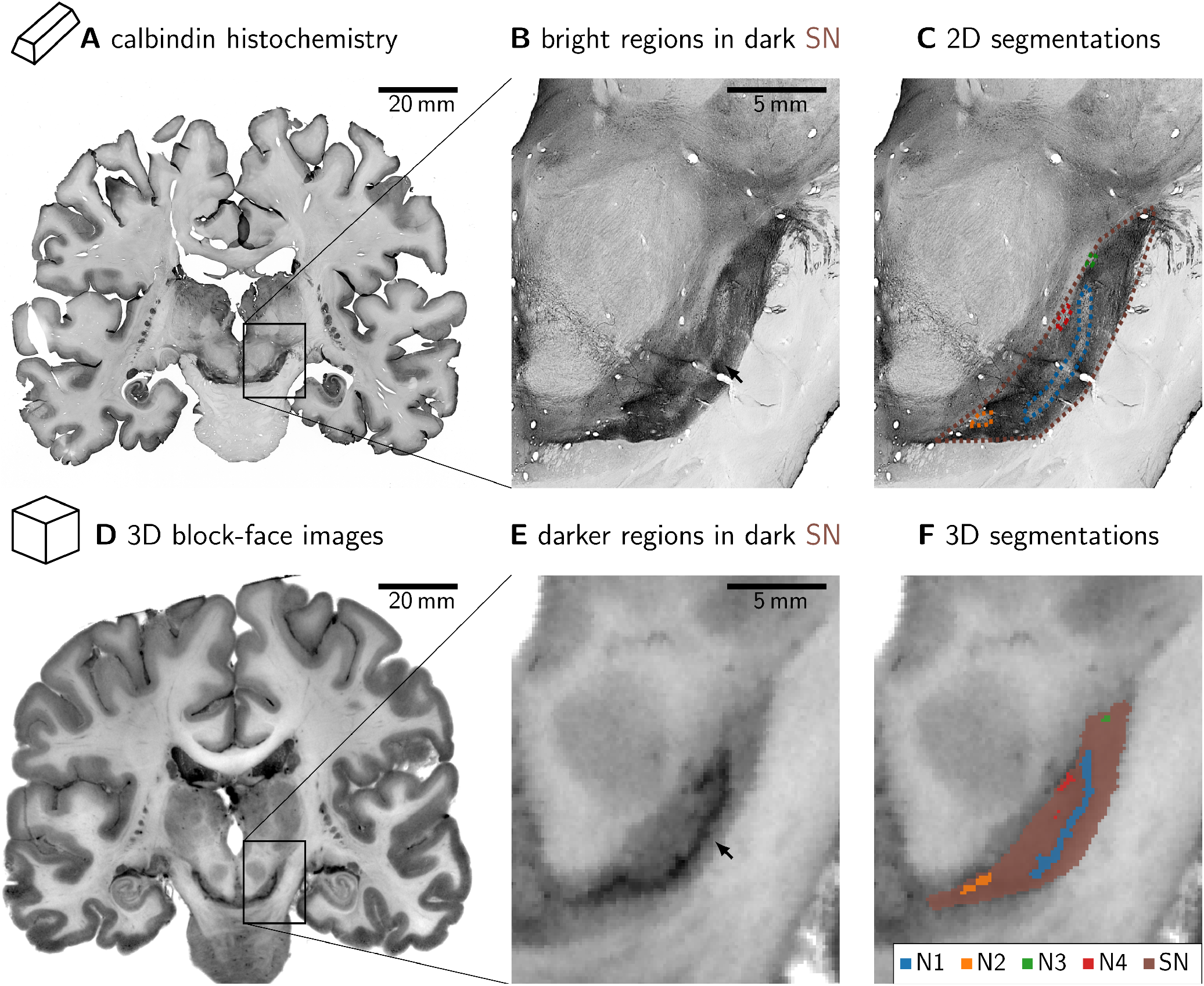
Gold-standard anti-calbindin immunohistochemistry validates 3D ultra-high-resolution nigrosome 1 (N1) and *substantia nigra* (SN) segmentations in 3D block-face images (BFI) of *post mortem* specimens. A, B, C: The nigrosomes were identified as areas of low anti-calbindin immunoreaction in two *post mortem* brain specimens (regions indicated with dotted colored outlines in C). A black arrow in B indicates the stripe-like appearance of N1. The SN was identified as a region of increased calbindin immunoreactivity. D, E, F: The nigrosomes were identified and segmented in 3D in gray-valued BFI as dark areas with high DN density and morphology in line with Damier *et al*.^3,5^. As in B, a black arrow in E indicates N1. The SN was segmented as an area darker than the surrounding tissue. We observed a high correspondence of nigrosome and SN segmentations in the BFI (F) and calbindin immunohistochemistry (C). C was manually affinely registered to F for easier comparison.

### Anti-calbindin-D28K immunohistochemistry

To verify the nigrosome and *substantia nigra* delineation, we used 200 µm sections stained with an antibody directed against calbindin-D28K^1^ at an 1:6 interval (donor 3 and 4), as described previously^22^ (Fig. 3D-3F). After thawing and rinsing of the sections, they were incubated overnight in the primary antibody. A biotinylated secondary antibody was used, and the signal was amplified using an avidine-biotinylated complex. The signal was visualised using 3,3’-diaminobenzidine as a chromogen. After mounting, dehydration, and coverslipping using Entellan, sections were imaged at a 21 µm in-plane resolution. The resulting images were co-registered to the BFI using a forward-backward approach across slices^19^, employing the ANTs SyN algorithm^23^.

### Histological gold-standard segmentations

To obtain gold-standard definitions of the *substantia nigra* and the nigrosomes, we used 3D-reconstructed anti-calbindin immunohistochemistry obtained on two *post mortem* tissue specimens described in the previous section, following the nigrosome and SN definitions of Damier *et al*.^3^ (Figs. 3A-3C). Calbindin staining was available at a 1:6 interval. All available calbindin sections of the SN were included in our analyses. We segmented the nigrosomes and the SN solely in those sections of the 3D-reconstructed images that corresponded to calbindin-stained sections. All segmentations were performed using the 3D Slicer^24^ and FSLeyes^25^ software.

#### Substantia nigra

Different histochemical techniques have been proposed to provide gold-standard outlines of the SN, which produce slightly different results^2^. These approaches are based on the expression of calbindin, enkephalin, and substance P^2^. For consistent SN segmentation on the sub-millimeter length scale, we used one of these definitions, based on the image contrast in 3D-reconstructed anti-calbindin microscopy images, to delineate the SN. In the area between the *nucleus ruber* and the *crus cerebri*, we found an area of increased immunohistochemical reactivity in the anti-calbindin microscopy images, which we defined as the SN.

#### Nigrosomes

We delineated the nigrosomes in 2D histological sections on which anti-calbindin immunohistochemistry had been performed, following their original definition^3^.

### Ultra-high-resolution 3D delineations

We created ultra-high-resolution 3D delineations of the *substantia nigra* and the nigrosomes using the image contrast in 3D block-face images (Figs. 3D-3F).

#### Nigrosomes

Areas with a high density of neuromelanin-pigmented DN and the morphology of the nigrosomes^3^ were segmented on BFI. A neuroanatomy expert (M.M.) segmented NM-rich areas in every tenth coronal section from posterior to anterior and instructed two non-expert raters in segmentation (C.Jan. and M.B.). These segmentation were subsequently filled and refined by one rater for each case and reviewed by the neuroanatomy expert. The procedure was iterated multiple times to ensure a high-quality segmentation, accurately reflecting the spatial arrangement of the nigrosomes^3^. We defined these DN-rich regions as nigrosomes.

#### Substantia nigra

In 3D BFI, the SN was delineated as a hypointense structure enveloped rostromedially by the *nucleus ruber* and ventrolaterally by the *crus cerebri*. In the posterior portion of the SN, the ventro-lateral border was sharp except for the most medial portion, which reflects a sharp increase in myelin content in the *crus cerebri*. There, we followed the bending of the hypointense, pigmented arc of nigrosome 1 (N1) and N2 (Fig. 3E), using N2 to define the medial border of the SN. The rostromedial border was more diffuse. For a consistent definition of this border, we followed the line between the rostromedial borders of N2 and N4. This resulted in a consistent segmentation in the posterior portion of the SN where the nigrosomes are situated. In the anterior portion of the SN, it was more challenging to define consistent, sharp borders in BFI and calbindin. We used the mamillary body as an anatomical reference for the SN’s consistent maximum anterior extent.

### Segmentation validation

To validate the 3D nigrosome and *substantia nigra* delineations, we compared them to the gold-standard calbindin immunohistochemistry in two specimens. To validate this 3D ultra-high-resolution SN delineation, we quantitatively compared it to the histological gold standard SN delineation described above. We calculated the Dice coefficients of both delineations on the sections in the BFI on which calbindin immunohistochemistry was available.

### Multimodal registration for creating the nigrosome atlas

We employed automated registration techniques to co-align the multi-modal histological and MRI data used in this study. First, we registered four *post mortem* datasets to a common space using *substantia nigra* delineations and created probability maps of the five nigrosomes. Second, we registered these nigrosome probability maps to the AHEAD template^6^ in the MNI152 2009b space^20^ to create a neuroimaging atlas of the nigrosomes. Third, to validate the nigrosome atlas, we registered an independent BigBrain dataset to the AHEAD template and compared our atlas of nigrosome 1 to it’s established contrast on *post mortem* MRI. Fourth, to demonstrate the application of our atlas, we registered the *in vivo* quantitative MRI dataset to the AHEAD template. For all registrations, we relied on the SyN algorithm in ANTs^23^, embedded in the ANTsPy^26^ and Nighres^27^ toolboxes.

#### Registration of nigrosome segmentations to create nigrosome probability maps

To bring the nigrosome segmentations to a common space, we co-registered the ultra-high-resolution 3D SN delineations of the four individual datasets to a common space, which was defined by the block face image of donor 3 (specimen # 7 in^22^). To this end, we applied a two-step registration procedure using AntsPy. We first used a smaller, then a larger gradient step of 0.1 and 0.2 as defined in AntsPy, respectively, to perform registrations with high and low regularization, respectively. We used the mean squared difference as a metric and the AntsPy default values of all other parameters. This registration enabled us to transform the nigrosome delineations of four cases into a common space, using nearest-neighbor interpolation. In this space, we created probability maps for each of the five nigrosomes by summing the binary nigrosome segmentations of the four cases and dividing the maps by the number of specimens. We applied the same procedure to the ultra-high-resolution 3D SN segmentations to obtain an SN probability map.

#### Registration of nigrosome probability maps to AHEAD template to create nigrosome atlas

To bring the nigrosome and SN probability maps to the standard MNI152 2009b space^20^, we co-registered the block-face image with the best image quality (specimen 3) to the multi-contrast quantitative MRI AHEAD template^6^ using the `focused_antsreg` function in Nighres^27^ (Fig. 4A). This function comprises a two-step ANTs registration procedure: First, it aligns whole-brain images and, second, focuses the registration around a given region. We registered the BFI to the quantitative proton density (*PD*), susceptibility (*χ*), *R*_1_ = 1*/T*_1_, where *T*_1_ is the longitudinal relaxation time, and 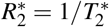 of the AHEAD template using cross-correlation as a registration metric (Figs. 4B-E). We first registered with high then with low regularization. We used the SN mask of the MASSP atlas^12^ for creating a registration focus with smoothly decreasing intensity outside of the SN mask. We applied the resulting transformation to the 3D ultra-high-resolution nigrosome and SN probability maps, yielding histological nigrosome and SN atlases in MNI152 2009b space.

**Figure 4.**
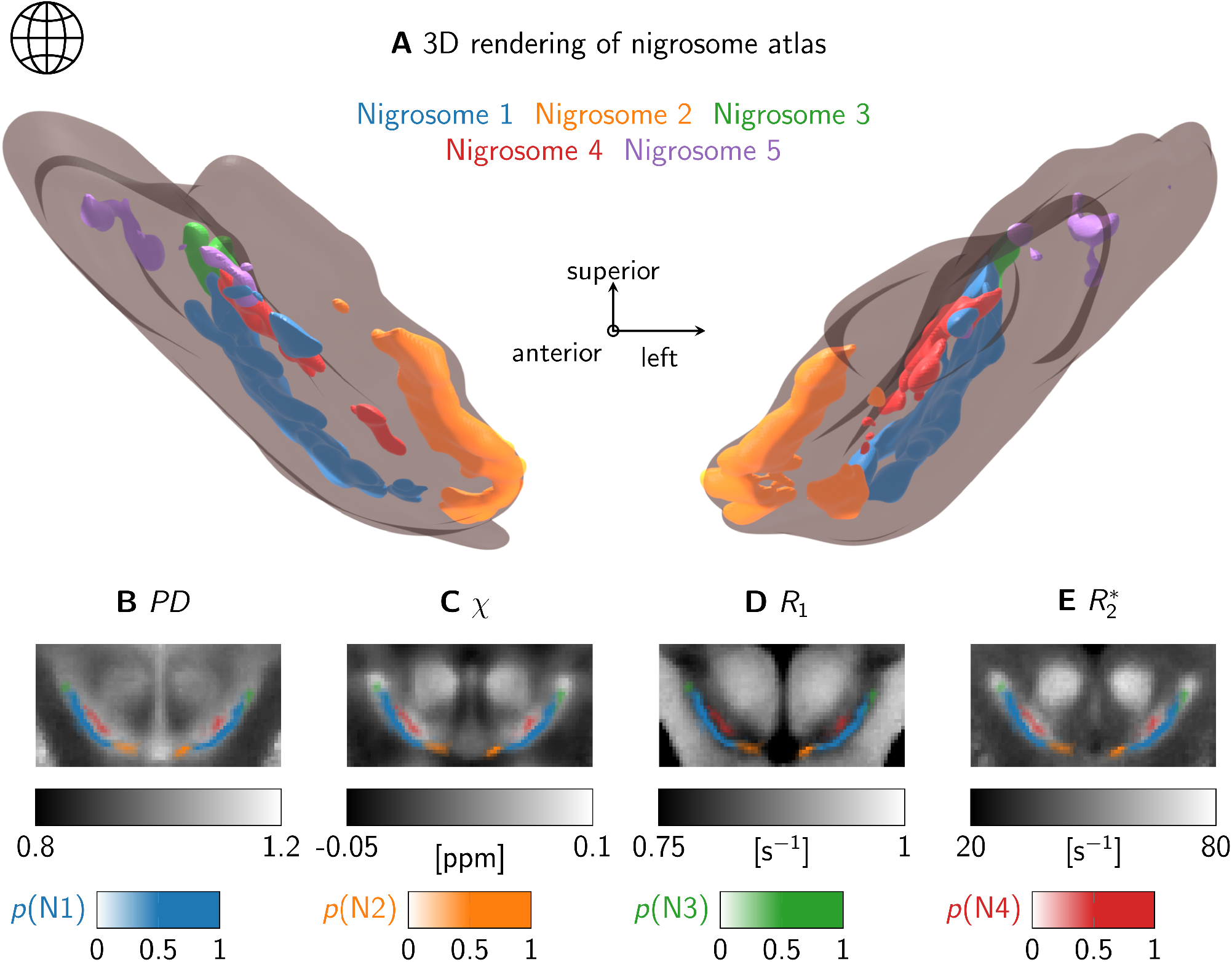
The nigrosome atlas aligned to the AHEAD template^6^ in MNI152 2009b template space^20^. A: A three-dimensional representation of the nigrosome atlas (colored volumes) within the *substantia nigra* (transparent gray volume). The view follows the anterior-posterior axis through the brain. For an accessible 3D view, we included a video of the nigrosome atlas in the data repository (https://osf.io/gsphy/files/nzxp2?view_only=ba16c0c39576450182181bfb8512317f). The anatomical reference frame is indicated, where the anterior direction points toward the viewer. B-E: The probability maps (*p*) of the nigrosomes 1-4 (N1-N4) are shown overlaid on the proton density (*PD*), susceptibility (*χ*), longitudinal relaxation rate (*R*_1_), and effective transverse relaxation rate 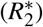 maps of the AHEAD template. As the nigrosomes do not extend over the entire SN, only nigrosomes N1-4 are visible in this section, which corresponds approximately to the histological section displayed in Fig. 1A. While some asymmetry of the probability maps is apparent, the positions of the nigrosomes and their spatial relationship to each other and the boundary between SN and the surrounding white matter are consistent with the original nigrosome definition (Fig. 1B). Moreover, N1 crosses and extends beyond the dorsolateral hypointensity in *χ* and 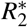 maps, as expected from previous research^10^.

### Nigrosome atlas validation against classical SN parcellations

To validate the resulting nigrosome atlas, we compared its topology with classical neuroanatomical descriptions by Halliday et al.^2^. We developed a method for automated parcellation of the three anatomical layers within the substantia nigra (SN). The probability maps of the five nigrosomes were projected onto these layers, and their relative positions were visualized in layer projections and compared with classical qualitative parcellations based on 2D histology.

#### Laplacian embedding of nigrosomes into layered SN representations

To define three discrete anatomical layers within SN corresponding to their dorsal tier, ventral tier and pars reticulata subdivisions we defined a coordinate system intrinsic to the SN with one axis aligned to SN’s thinnest dimension. A graph between all voxels inside the SN was created with distances defined as a weighted combination of Euclidean distance and distance to the SN boundary as in^28^, using the `spectral_voxel_thickness_embedding` function in Nighres^27^. The obtained graph was transformed by approximate Laplacian embedding^29^, where the first three dimensions of embedding define a curved coordinate system following the shape of the SN (Fig. 5). The embedding across the thickness of the SN was subdivided into three layers approximating the anatomically defined tiers, based on histograms of the nigrosome relative depths (Fig. 5B), and the probabilistic maps of each nigrosome were averaged inside each layer and displayed along the two orthogonal embedding directions (Fig. 5C).

**Figure 5.**
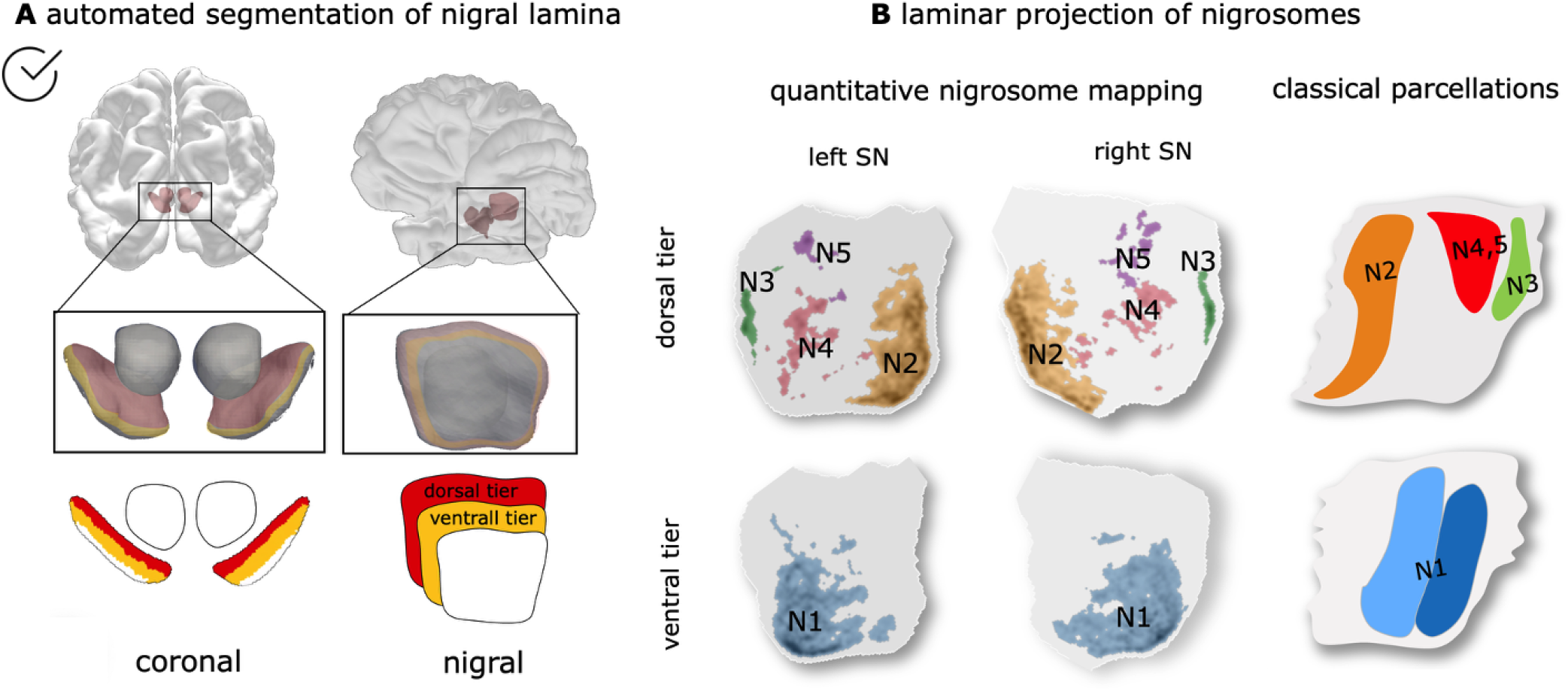
Validation of the anatomical plausibility of the 3D nigrosome atlas against classical *substantia nigra* (SN) delineations. A: Top: Translation of the classical delineations into the space of the 3D atlas. Anatomical layers within the *substantia nigra* (SN) were defined based on shape analysis. The thinnest dimension was automatically determined, and the SN was subdivided into three layers approximating the ventral and dorsal tiers (*pars compacta*) and *pars reticulata*. Bottom: Automated segmentation of the SN into three layers based on geometric analysis in a coronal slice. B: Left: Projection of the 3D nigrosome atlas onto the defined layers. Right: Arrangement of nigrosomes within the dorsal tier (nigrosome 1) and ventral tier (nigrosomes 2–5). The schematic was adapted from^2^. The relative spatial arrangement of the nigrosomes in our atlas corresponds well to qualitative histological delineations, demonstrating the anatomical plausibility of the 3D atlas.

### Independent multi-modal dataset for technical validation

To assess the accuracy of the automated registration procedure ultra-high resolution nigrosome atlas, we employed an independent dataset comprising ultra-high-field and -resolution *post mortem* MRI ^30^ (Fig. 6). Here, we outline tissue processing and MRI acquisition.

**Figure 6.**
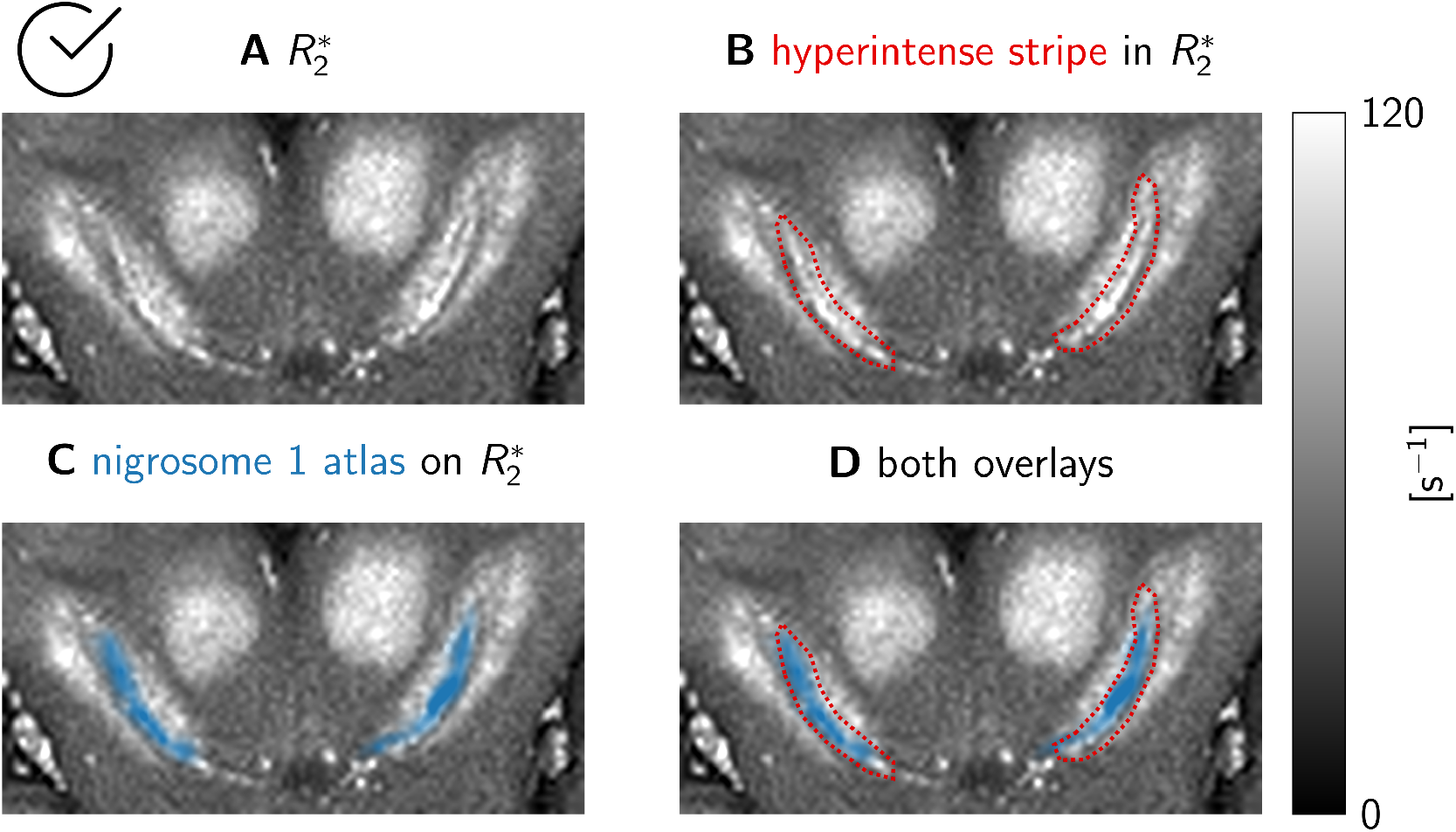
Validating the nigrosome atlas using the BigBrain dataset^21^. To validate the accuracy of our nigrosome atlas, we reproduce a finding that we previously reported: nigrosome 1 appears as a hyperintense stripe in 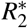 maps of *post mortem* brain specimens^10^. In an 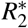 map acquired on the BigBrain specimen (A), a hyperintense stripe is visible in the *substantia nigra* (marked in red in B). After using the registration routine, we propose to apply our nigrosome atlas to align the latter to the BigBrain dataset. Nigrosome 1 (blue area in C) overlaps strongly with the hyperintense stripe (overlay in D). This suggests that the proposed registration routine is precise enough to enable neuroimaging of the nigrosomes. Furthermore, the nigrosome anatomy captured by our atlas captures the anatomy in this independent dataset accurately.

#### Tissue processing

A *post mortem* human brain (female, age of death: 73 years, cause of death: acute respiratory syndrome, *post mortem* time before fixation: 7 hours) was fixed in 4 % paraformaldehyde solution for four months. Tissue acquisition, handling, and fixation procedures followed the procedure described in detail in^21^.

#### MRI

The brain was transferred to a phosphate-buffered solution (PBS, pH 7.4) 72 hours before scanning to remove formaldehyde monomers, thereby improving image contrast. The brain was covered and padded with cotton tissues for MRI scanning to prevent cortical damage. Then, it was placed in a custom-made head-shaped container filled with degassed PBS. The container was exposed to 1 mbar vacuum for 12 hours to remove air bubbles. Multi-parameteric mapping (MPM)^31^ data were acquired on a 7 T system (7 T Terra, Siemens Healthineers, Erlangen, Germany), using a 32-channel radio-frequency head coil (Nova Medical, Wilmington, USA). Four multi-echo 3D FLASH acquisitions were obtained with different weightings: proton-density-weighted, longitudinal-relaxation-time (*T*_1_)-weighted, magnetization-transfer (*MT*)-weighted, and a scan at Ernst angle. The following parameters were used: repetition time (*T*_*R*_)= 50 ms; eight equidistant echoes with echo times *T*_*E*1…8_ = 3.66 22.56 ms acquired using bipolar readout; isotropic resolution of 0.3 mm (field-of-view (*FOV*) = 192 mm, matrix 640 × 640); and a bandwidth of *BW* = 434 Hz*/*pixel. Excitation flip angles (*FA*) were *FA* = 11^°^ for the proton density and magnetization transfer-weighted scans, *FA* = 25^°^ for the Ernst angle scan, and *FA* = 59^°^ for the *T*_1_-weighted scan. An additional calibration scan for *FA* mapping was obtained using the Bloch-Siegert-Shift method^32^. The quantitative maps 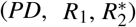 were reconstructed using a customized version of the hMRI toolbox (hMRI.info), adapted for *post mortem* and 7 T imaging^33^, using a three-flip-angle calculation without applying a small-flip-angle approximation.

#### Registration of nigrosome atlas to independent *post mortem* dataset

To validate the nigrosome atlas, we mapped it onto an independent, newly acquired BigBrain dataset that was histologically processed according to^21^. This dataset comprises quantitative maps of *R*_1_ and 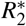, and a semi-quantitative *PD* map, all with 0.3 mm isotropic resolution.

Employing the `focused_antspy` function in Nighres as above, we registered these maps to the corresponding maps of the AHEAD template. We used the registration result to transform the nigrosome atlas to the space of the BigBrain dataset.

### *In vivo* quantitative MRI for usage illustration

To demonstrate the usage of the nigrosome atlas, we aligned it to *in vivo* MRI datasets acquired on four healthy volunteers, using the registration procedure described above (Fig. 7).

**Figure 7.**
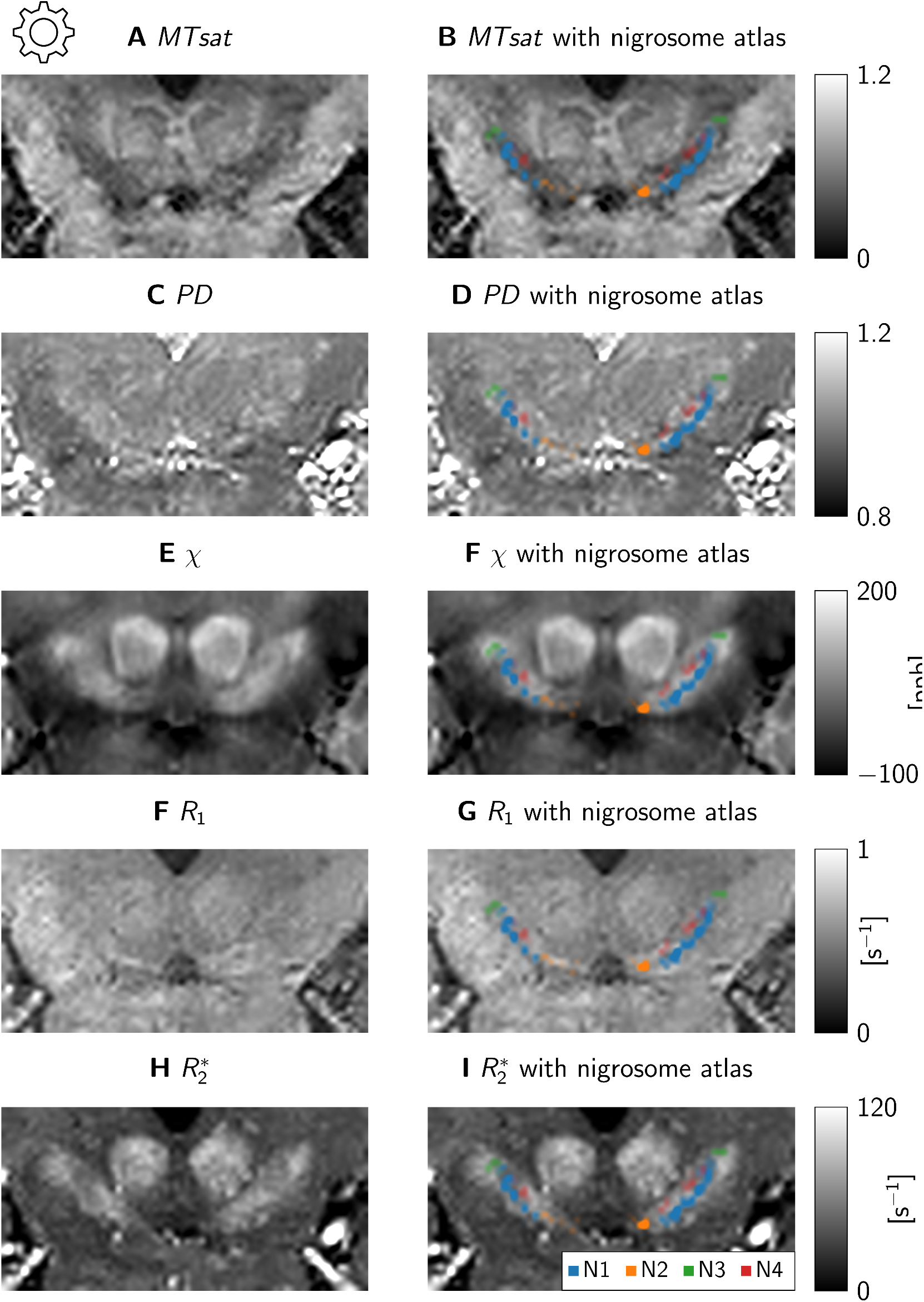
A usage example of the nigrosome atlas. Using the proposed dedicated registration, we aligned the nigrosome atlas to four *in vivo* quantitative MRI datasets, of which one representative example is shown. The dataset comprises quantitative maps of magnetization transfer saturation *MTsat* (A, B), proton density *pd* (C, D), longitudinal relaxation rate *R*_1_ (E, F), and effective transverse relaxation 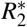 (G, H). The accuracy of the nigrosome atlas and the used registration is underscored by the alignment of N1 with the border between the hypointense *substantia nigra* and hyperintense *crus cerebri* in the *MTsat* map: N1 is located precisely on the border but inside the *substantia nigra*. Note that the hyperintense region encompassing the *substantia nigra* in the 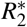 map shows a more lateral border to *crus cerebri*, as has been reported before^44^.

#### MRI

To obtain this dataset, four healthy adult participants (1 female, mean age (45.25 ± 6.30) years), each took part in at least two scanning sessions on a 7 T system (7 T Terra, Siemens Healthineers, Erlangen, Germany) using parallel transmission (pTx) and an 8 transmit-/32 receive-channel radiofrequency head coil (Nova Medical, Wilmington, USA). Scanning was performed across two sites: MPI CBS in Leipzig, Germany, and GIGA-Institute in Liège, Belgium.

MPM acquisition consisted of three whole-brain multi-echo 3D FLASH acquisitions with different weightings (*PD, T*_1_ and *MT* -weighted) with parallel transmit (pTx) kT-points excitation at an isotropic resolution of 0.6 mm (*FOV* = 218 mm × 250 mm × 173 mm, matrix = 364 × 416 × 288). The following parameters were used: *T*_*R*_ = 22.4 ms, *FA* for *PD*w/*MT* w/*T*_1_w were 7^°^*/*7^°^*/*22^°^, with six (four for MT-weighted) equidistant echoes *T*_*E*1…6_ = 3.00 15.6 ms. Parallel imaging using CAIPIRINHA with an acceleration factor *R* = 2 × 2 enabled acquisition time 8.25 min per contrast. *B*_1_ mapping for transmit field correction was performed using the AFI method with kT-points excitation^34^. The *B*_1_ mapping was repeated with a conventional excitation pulse to correct for spatial inhomogeneity in the MT saturation. The total acquisition time, including shimming, was around 35 min.

Images were reconstructed with AC-LORAKS^35,36^ and MCPC-3D-S^37^ followed by LCPCA denoising^12^. The images were processed with the open source hMRI toolbox^33^, yielding 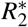, *PD, MTsat*, and *R*_1_ maps. Subcortical structures were automatically parcellated by MASSP using all three contrasts as input^12^. QSM was additionally calculated for each contrast by QSMxT^38^ and combined using the robust combination feature of the hMRI toolbox^39^, weighting the contrasts by the inverse of the quality maps produced by QSMxT’s phase-unwrapping step, Rapid open-source minimum spanning tree algorithm (ROMEO)^40^.

#### Registration nigrosome atlas to *in vivo* dataset

To demonstrate the usage of the nigrosome atlas, we applied it to an *in vivo* MRI dataset comprising quantitative maps of proton density, *R*_1_, and 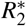 with an isotropic resolution of 0.6 mm.

To align the *in vivo* MRI data to the AHEAD template, we employed the `focused_antspy` function in Nighres as above, registering the *in vivo* quantitative maps to the corresponding map of the AHEAD template. We used the registration result to transform the nigrosome atlas to the space of the *in vivo* MRI dataset.

## Data Records

The data repository^41^ (https://osf.io/gsphy/overview?view_only=ba16c0c39576450182181bfb8512317f) contains four folders: *probability_maps* contains the nigrosome atlas, *scripts* contains registration scripts to apply the atlas, *singularity* contains a singularity container to run the registration scripts reproducibly, and *ahead_template* contains the AHEAD template needed in the registration scripts. The repository contains a README file that explains the data structure and usage of the repository. Furthermore, a video (https://osf.io/gsphy/files/nzxp2?view_only=ba16c0c39576450182181bfb8512317f) in the repository shows a 3D rendering of the nigrosomes.

In the folder *probability_maps*, the nigrosome atlas is provided as a collection of probability maps of the five nigrosomes in the MNI152 2009b Nonlinear Asymmetric template space^20^, aligned to the AHEAD dataset^22^ at 0.5 mm isotropic resolution. Together with the nigrosome atlas, we provide a *substantia nigra* probability map based on block-face images. We provide all maps in a compressed NIfTI-1 data format (…*nii*.*gz*). Fig. 4 shows a 3D rendering of the probability maps and those maps overlayed on the quantitative MRI parameter maps of the AHEAD template.

The other folders contain all the data needed to apply the atlas. The *scripts* contains a Jupyter notebook script that registers the nigrosome atlas to MRI data as described above. For ease of use, in the *singularity* folder, we provide a Singularity container with all the necessary software to run the registration scripts. To enable a standalone execution of the registration scripts, the AHEAD template^22^ is provided in the *ahead_template* folder.

## Technical Validation

In the following, we describe three validation procedures of the nigrosome atlas: the validation of the anatomical delineations, the plausibility of 3D nigral atlas in light of classical nigral anatomical delineations, and the validation of the registration accuracy.

### Segmentation validation

First, we validated the nigrosome and *substantia nigra* segmentations in 3D histology using gold standard calbindin immunohistochemistry (Fig. 3). We qualitatively confirmed the low anti-calbindin immunoreactivity of the 3D nigrosome segmentations in two specimens: We assessed that corresponding coronal sections of BFI and calbindin immunohistochemistry showed the same subset of nigrosomes with a similar spatial arrangement. We note that a quantitative comparison, which would have been preferable, was unfeasible due to the registration precision between the BFI and the calbindin immunohistochemistry. The quality of this registration is impeded by severe non-linear deformations of histological sections floating in a liquid after cutting. This resulted in a registration accuracy lower than the extent of the nigrosomes (≈0.5 mm). This rendered a quantitative comparison impossible. In 3D, the nigrosome segmentations appeared as partly disconnected volumes, reflecting the reticular buildup of the dopaminergic neurons in the SN^42^ (Fig. 4, video in data repository).

To validate the *substantia nigra* segmentations based on BFI and used for registering the nigrosome segmentations, we quantitatively compared them to SN segmentations on calbindin immunohistochemistry. The Dice coefficient of the SN segmentations based on BFI and calbindin immunohistochemistry were 86 % and 79 % for the two specimens, underscoring a high accuracy of the SN segmentation in BFI.

### Anatomical plausibility

To assess whether the complete procedure of nigrosome segmentation and inter-subject co-alignment preserves the anatomical principles proposed for nigrosome organization, we first qualitatively compared the automated layering of the substantia nigra (SN, Fig. 5C) with classical delineations of the dorsal tier, ventral tier, and pars reticulata (Fig. 5A). A high degree of similarity was observed between layers segmented using Laplacian embedding and classical anatomical delineations (Fig. 5). We then compared the relative arrangement of nigrosomes within the dorsal and ventral tiers with classical descriptions. A strong correspondence was observed between our fully three-dimensional, data-driven approach and classical delineations, with nigrosome 1 predominantly located in the medial portion of the ventral tier, and nigrosomes N2, N3, N4, and N5 arranged within the dorsal tier (compare bottom raw Fig. 5A, Fig. 5C). This close correspondence between established principles of SN organization and our atlas highlights its anatomical plausibility.

### Registration validation

To demonstrate the accuracy of the nigrosome atlas, we applied it to the independent *post mortem* dataset (Fig. 6). This dataset comprises 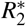 maps, which show a hyperintense stripe within the SN (Fig. 6A, 6B). In earlier work, we have shown that this stripe corresponds to nigrosome 1^10^. After using the registration procedure we proposed to align the nigrosome atlas to this quantitative MRI dataset, we observed a high correspondence of the nigrosome 1 atlas to the hyperintense stripe (Fig. 6C, 6D). Note that the qMRI maps of the AHEAD template used for registration do not show nigrosomes as areas of increased 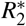, indicating that using the SN contrast for registration provides enough accuracy to align the nigrosomes contained in the SN. This demonstrates that the registration procedure we proposed for applying the nigrosome atlas achieves a high enough accuracy for nigrosome imaging.

## Usage Notes

We provide a histology-based 3D atlas of the five nigrosomes in the *substantia nigra* together with scripts that facilitate its registration to MRI data recorded at 7 T. The registration relies on a focused multi-contrast registration provided by the Nighres software package.

To demonstrate the use of our atlas, we applied it to a dataset of quantitative multi-parameteric mapping MRI data recorded at 7 T using the focused registration as in the provided script. The registration relies on a two-step alignment of quantitative MRI maps to the AHEAD template maps, comprising *PD, χ, R*_1_, and 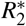. After a brain-wide alignment in the first step, the second step focuses on the SN region defined by our atlas, thereby improving the registration accuracy in this region. The registration to the AHEAD dataset can be done by using only a subset of the provided quantitative MRI maps, which may have a negative impact on registration accuracy. Thus, if fewer quantitative MRI maps are available, a critical evaluation of the registration accuracy is indispensable.

Note that although published in MNI152 2009b asymmetric space, the AHEAD template differs significantly from the original MNI152 2009b template on the nigrosome length scale, hence we advise against using the MNI152 2009b to align our atlas to MRI data.

## Code availability

Code for aligning the nigrosome atlas is available in the data repository as a Jupyter notebook^41^. As a use case, we applied the nigrosome atlas to an *in vivo* quantitative MRI dataset using the provided registration script (Fig. 7).

All code for generating the nigrosome atlas can be provided upon reasonable request.

## Acknowledgements

We thank Patrick Scheibe for his help with data visualization. We express our gratitude to Luke J. Edwards and Niklas Kügler for their help with processing MRI data, and to Sara Schaumberg for her help with segmenting histological data. We thank the whole-body donors and acknowledge the whole-body donation program at the University of Maastricht for providing *post mortem* brain samples. This is an EU Joint Programme - Neurodegenerative Disease Research (JPND) project. The project is supported through the following funding organisations under the aegis of JPND - www.jpnd.eu: the German Federal Ministry of Education and Research (BMBF) under grant numbers 01ED2210 and 01ED2508, the Dutch Organisation for Knowledge and Innovation in Health, Healthcare and Well-being (ZonMw) under grant numbers 10510062420003 and 10510062110003.

## Author contributions statement

Conceptualization: MB, AA, PLB, NW, EK; Methodology: MB, AA, PLB, CJan, CJäg, AH, KJP, MM, RB, EK; Software: MB, PLB; Validation: MB, AA, PLB, EK; Formal analysis: MB, PLB, MM, EK; Investigation: MB, AA, PLB, MM, EK; Resources: AA, KA, BUF, NW, EK; Data Curation: MB; Writing - Original Draft: MB, PLB, EK; Writing - Review & Editing: MB, AA, PLB, CJan, CJäg, AH, KJP, MM, RB, KA, BUF, NW, EK; Visualization: MB, PLB, EK; Supervision: MB, PLB, MM, NW, EK; Project administration: MB, NW, EK; Funding acquisition: MB, AA, MM, KA, BUF, NW, EK;

## Footnotes

1 In the rest of the manuscript, we will omit the D28K for readibility and refer to calbindin, which is always calbindin-D28K.

